# HiC-DC+: systematic 3D interaction calls and differential analysis for Hi-C and HiChIP

**DOI:** 10.1101/2020.10.11.335273

**Authors:** Merve Sahin, Wilfred Wong, Yingqian Zhan, Kinsey Van Deynze, Richard Koche, Christina S. Leslie

## Abstract

We present HiC-DC+, a software tool for Hi-C/Hi-ChIP interaction calling and differential analysis using an efficient implementation of the HiC-DC statistical framework. HiC-DC+ integrates with popular preprocessing and visualization tools, includes TAD and A/B compartment callers, and outperformed existing tools in H3K27ac HiChIP benchmarking as validated by CRISPRi-FlowFISH. Differential HiC-DC+ analysis recovered global principles of 3D organization during cohesin perturbation and differentiation, including TAD aggregation, enhancer hubs, and promoter-enhancer loop dynamics.

## Main

While there are effective software tools for visualizing 3D genomic interaction data from Hi-C and related assays, existing statistical analysis tools for these data are less mature, and integrative genome-wide analyses remain challenging. The HiC-DC+ (Hi-C/HiChIP direct caller plus) package enables principled statistical analysis of Hi-C and HiChIP data sets – including calling significant interactions within a single experiment and performing differential analysis between conditions given replicate experiments – to facilitate global integrative studies (**Fig. 1a**).

**Figure 1.**
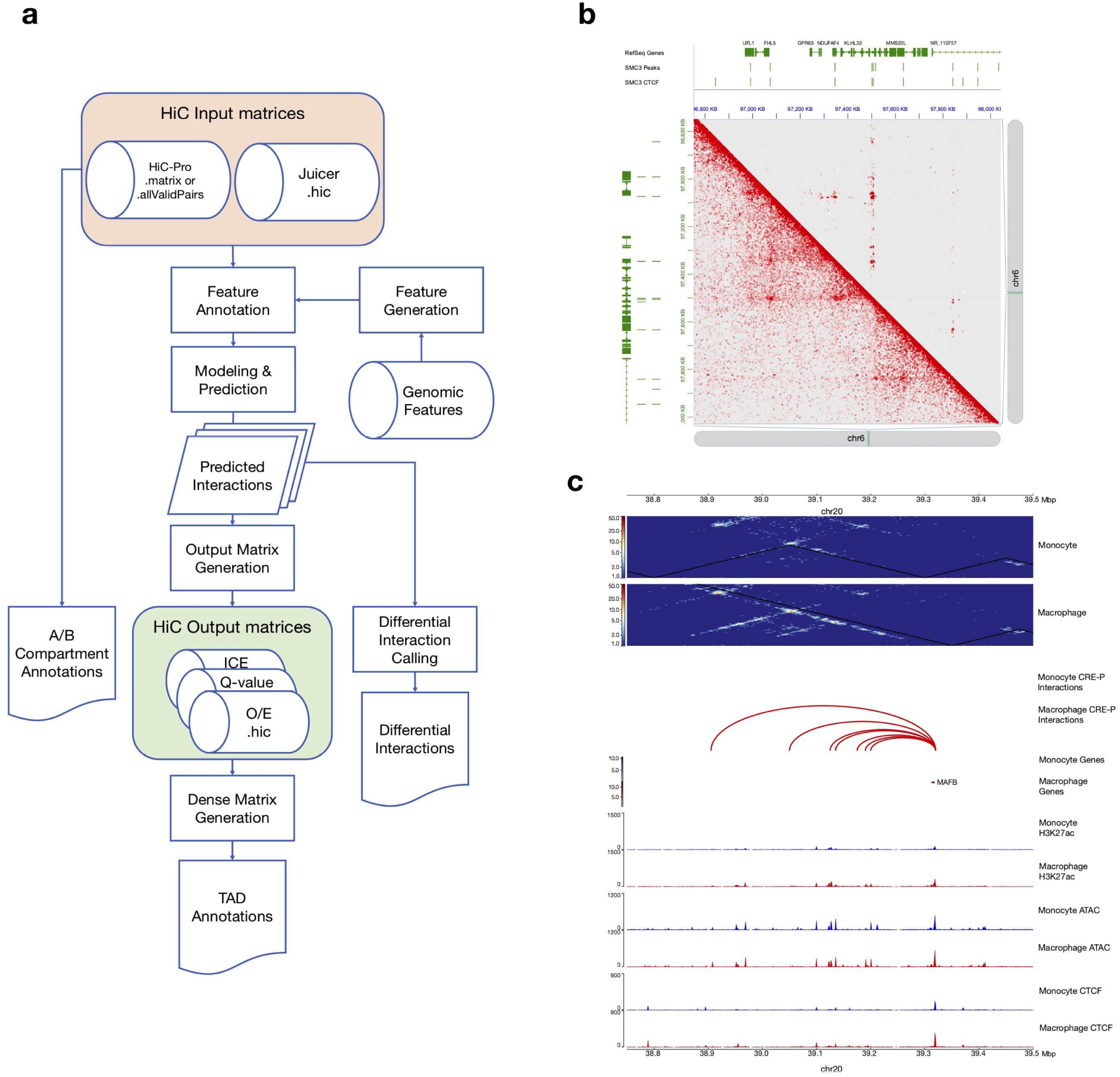
HiC-DC+ workflow and capabilities. **a.** Overview of HiC-DC+ pipeline. **b.** Visualization of HiC-DC+ results in Juicer: FDR-corrected *P* values shown in the upper triangle heatmap, KR normalized counts in the bottom triangle, for SMC1a HiChIP in GM12878 at 5kb resolution. **c**. Visualization of HiC-DC+ differential interactions in HiCExplorer: HiC-DC+ Z-score normalized Hi-C counts in THP-1 monocytes and macrophages at 5kb resolution are shown as heatmaps, along with differential Hi-C interactions as arcs and ChIP-seq and ATAC-seq as signal tracks for a region encompassing the gene *MAFB*.

HiC-DC+ estimates significant interactions in a Hi-C or HiChIP experiment directly from the raw contact matrix for each chromosome up to a specified genomic distance, binned by uniform genomic intervals or restriction enzyme fragments, by training a background model to account for random polymer ligation and systematic sources of read count variation. Similar to HiC-DC^1^, HiC-DC+ uses negative binomial regression to estimate the expected read count in an interaction bin based on genomic distance, and the GC content, mappability, and effective bin size based on restriction enzyme (RE) sites in the corresponding pair of genomic intervals (**Methods**); genomic features derived from the set of RE recognition sites are provided as input to the model. As in HiC-DC and Fit-Hi-C^2^, HiC-DC+ uses a two-step model fitting procedure to increase power (**Methods**). Given the higher coverage of current Hi-C and HiChIP experiments, HiC-DC+ does not incorporate zero truncation in the background model, resulting in a somewhat simpler statistical model than used by HiC-DC.

HiC-DC+ also enables differential analysis of Hi-C or HiChIP interactions between a pair of conditions, given replicate experiments. To do this, HiC-DC+ estimates genomic distance-dependent scaling factors from the data and uses DESeq2 to assess the significance of differential interactions (**Methods**). This approach is similar to the ACCOST method^3^ but exploits existing capabilities of DESeq2 to perform the statistical test.

The HiC-DC+ Bioconductor R package represents a major software update relative to HiC-DC, with an efficient implementation of the core statistical model, data files and storage objects and the ability to parallelize. These features are required to process very deep (≥ 2B reads) data sets at high resolution. The package supports standard Hi-C contact matrix file formats (.hic, .matrix, .allValidPairs) generated by HiC-Pro or Juicer as input (**Fig. 1a**). For convenience, the HiC-DC+ package also implements callers for topologically associating domains (TADs), through an implementation of TopDom, and A/B compartments. The TAD caller can be run on contact matrices that have been normalized by HiC-DC+, namely observed/expected (O/E) counts or negative binomial Z-score values (**Fig. 1a**, **Methods**), or by other methods such as ICE^4^ .

To visualize significant HiC-DC+ 3D interactions and differential loops, results from the package (FDR-corrected *P* values, or O/E or Z-score normalized counts in .hic file format) can be readily supplied to popular visualization tools such as Juicer^5^ and HiCExplorer^6^. For example, **Fig. 1b** shows the results of HiC-DC+ analysis of a SMC1a HiChIP data set in the GM12878 lymphoblastoid cell line^7, 8^ in a 1.2Mb region of chr 6 using Juicer, with KR normalized count data below the diagonal and significant interactions at FDR < 0.05 above the diagonal. HiCExplorer is useful for visualizing HiC-DC+ normalized contact matrices in triangular format and significant interactions as arcs, alongside other genomic tracks. In **Fig. 1c**, HiC-DC+ Z-score normalized Hi-C data in untreated and PMA-treated THP-1 cells^9^ are shown at a 1Mb region encompassing *MAFB*, an important regulator of monocyte to macrophage differentiation, together with HiC-DC+ differentially gained enhancer-promoter interactions and signal tracks for ATAC-seq, H3K27ac, and CTCF ChIP-seq.

HiChIP enriches for 3D interactions associated with a protein or histone modification of interest by applying ChIP to the contact library after nuclear lysis^8^. The HiC-DC+ statistical model proved to be well suited for HiChIP analysis without requiring any additional covariates, such as signal from a parallel ChIP-seq experiment or HiChIP self-ligation read counts, which have been used to mimic the ChIP-seq signal^8^. By contrast, some existing HiChIP analysis methods like hichipper^10^ and FitHiChIP^11^ indeed attempt to call genomic peaks directly from HiChIP or use a parallel ChIP-seq experiment as a step in their modeling.

To benchmark HiC-DC+ on HiChIP, we ran the model on existing H3K27ac HiChIP in K562 cells^8^ at 5kb resolution and assessed using recently generated CRISPRi-FlowFISH data for 22 genes^12^ (**Methods**). CRISPRi-FlowFISH quantifies enhancer function in terms of the estimated log fold change of target gene expression due to enhancer inactivation by using a pooled CRISPRi screen against candidate enhancers and sorting cells using RNA fluorescence in situ hybridization (FISH) against the target gene. We reasoned that a successful H3K27ac HiChIP analysis pipeline should identify significant loops between the target gene promoter and FlowFISH-validated accessible sites, and rank these loops above interactions between the target promoter and accessible sites with insignificant FlowFISH effect.

Therefore, for each CRISPRi-FlowFISH experiment and its corresponding target gene, we ranked candidate interactions at 5kb between the target promoter and each tested regulatory element by HiC-DC+ *P* value. Similarly, we ranked candidate interactions using three other published methods: Fit-HiChIP^11^, an adaptation of Fit-Hi-C for HiChIP; MAPS^13^, another approach based on fitting a generalized linear model to estimate a background distribution; and hichipper^10^, a method that models the background read density in terms of proximity to RE sites. When we evaluated performance per gene by auPR (area under the precision-recall curve), HiC-DC+ outperformed hichipper and MAPS (**Fig. 2a**, *P* < 0.01, signed rank test). Although per gene auPR was not significantly different between HiC-DC+ and FitHiChIP (*P* = 0.15, signed rank test), HiC-DC+ had a higher median per gene auPR (0.307 for HiC-DC+ vs 0.245 for FitHiChIP). Moreover, when we examined the predicted interactions at FDR < 0.01 across all genes for each method, we found that HiC-DC+ was most successful in identifying promoter-enhancer loops of enhancers with large effect size on target gene expression as assessed by CRISPRi-FlowFISH (**Fig. 2b,** *P* < 0.011 FitHiChIP, *P* < 0.005 for MAPS and *P* < 0.001 for hichipper, rank sum test). While Fit-HiChIP was the runner-up in terms of mean auPR (0.180 for FitHiChIP vs. 0.205 for HiC-DC+), suggesting a reasonable ranking of promoter-to-candidate-enhancer interactions, it called many more interactions than HiC-DC+ at a fixed FDR < 0.01 threshold, indicating that its *P* values are somewhat inflated. Indeed, at the PLP2 and FTL loci (**Fig. 2c,d**), FitHiChIP identifies both positive and negative enhancers but generates a large number of false positive interactions, whereas HiC-DC+ identifies significant interactions that coincide with positive enhancers and H3K27ac peaks.

**Figure 2.**
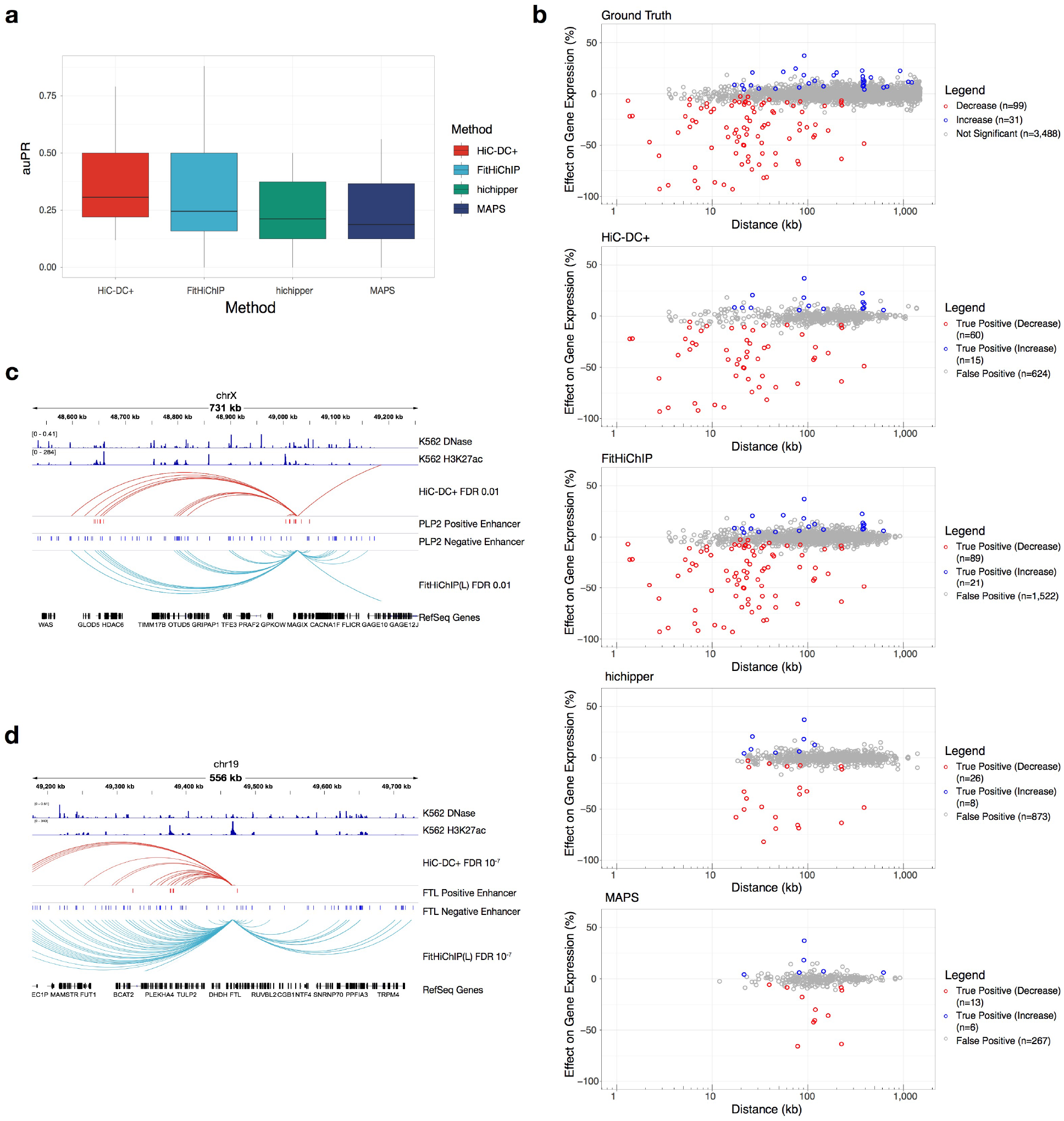
HiC-DC+ accurately identifies enhancer-promoter interactions from H3K27ac HiChIP. **a.** Evaluation of method performance by per gene auPR values. For each target gene, we ranked the candidate regulatory elements tested by CRISPRi-FlowFISH in K562 cells^12^ based on the significance (*P* value) or corresponding 5kb HiChIP interactions with the target promoter as estimated by each method. **b.** Comparison of performance for identifying promoter-enhancer loops with large effect sizes on target gene expression as assessed by CRISPRi-FlowFISH. Each dot in the scatterplots represents one tested promoter-enhancer pair: true positives resulting in decreased target gene expression upon enhancer inactivation are shown in red, true positives resulting in increased target gene expression shown in blue, false positives in grey. **c.** Promoter-enhancer loops identified by HiC-DC+ and FitHiChIP (FDR < 0.01) are shown as arcs for *PLP2* gene, along with all candidate enhancers tested by CRISPRi-FlowFISH^12^ and signal tracks for K562 DNase-seq and H3K27ac ChIP-seq. **d.** Promoter-enhancer loops identified by HiC-DC+ and FitHiChIP (FDR < 10^-7^) are shown as arcs for *FTL* gene, along with all candidate enhancers tested by CRISPRi-FlowFISH and K562 DNase-seq and H3K27ac ChIP-seq signal tracks.

Despite evidence that promoter-enhancer interactions identified by CRISPRi-FlowFISH are enriched for shorter range interactions that are detectable by H3K27ac signal and accessibility alone (**Supplementary Material; Supplementary Fig. 1**), enhancer screening data provides a useful benchmark data set for comparing HiChIP interaction callers.

As a second test case, we benchmarked HiC-DC+ and existing HiChIP interaction callers on published HiChIP data for SMC1a, a subunit of the cohesin complex, in GM12878^8^. These methods called vastly different numbers of significant cohesin-mediated loops at 5 kb resolution with genome-wide FDR < 0.01: 81,861 loops for HiC-DC+, 95,196 for Fit-HiChIP, 38,894 for MAPS, and 52,590 for hichipper. When we restricted to loops whose anchors both contained CTCF motifs, a majority of such loops (75%-84%) for all methods except hichipper (49%) were associated with a pair of convergently oriented motifs (**Supplementary Fig. 2**). We also examined the enrichment of significant interactions from each method at the boundaries of previously reported subTADs in GM12878 based on Hi-C analysis^7^. We found that HiC-DC+ interactions were most highly enriched at subTAD boundaries, with Fit-HiChIP as runner-up (**Supplementary Fig. 3**).

To call differential Hi-C and HiChIP interactions between a pair of conditions, we estimate distance dependent DESeq2 size factors so that the median normalized count for pairs of bins at each given distance in each matrix is the same (**Methods**). Dispersion and MA plots behave as expected, suggesting that our normalization works appropriately (**Supplementary Fig. 4**).

To benchmark our differential HiC interactions against diffHiC^14^, multiHiCcompare^15^, and Selfish^16^, we ran HiC-DC+ and other tools on Hi-C data in HAP1 and WAPL knockout HAP1 cells at 25 kb resolution^17^. Haarhuis et al. showed that removing WAPL affects chromosome topology on a global scale through the formation of longer loops and strongly increased interaction frequencies between nearby TADs^17^. In particular, they highlighted three genomic regions to show accumulation of contacts at TAD corners (**Fig. 3a**, top row) and loop extension linked to TAD aggregation (**Fig. 3a**, middle and bottom rows) upon WAPL loss. HiC-DC+ can detect these enriched loops more effectively than other methods at FDR < 0.05. Other methods also call at least two times more significant interactions than HiC-DC+, suggesting *P* value inflation (**Supplementary Fig. 5**). The loop lengths of differential interactions with HiC-DC+ span a wide range, while other methods appear biased towards short- or long-range interactions (**Supplementary Fig. 6**).

**Figure 3.**
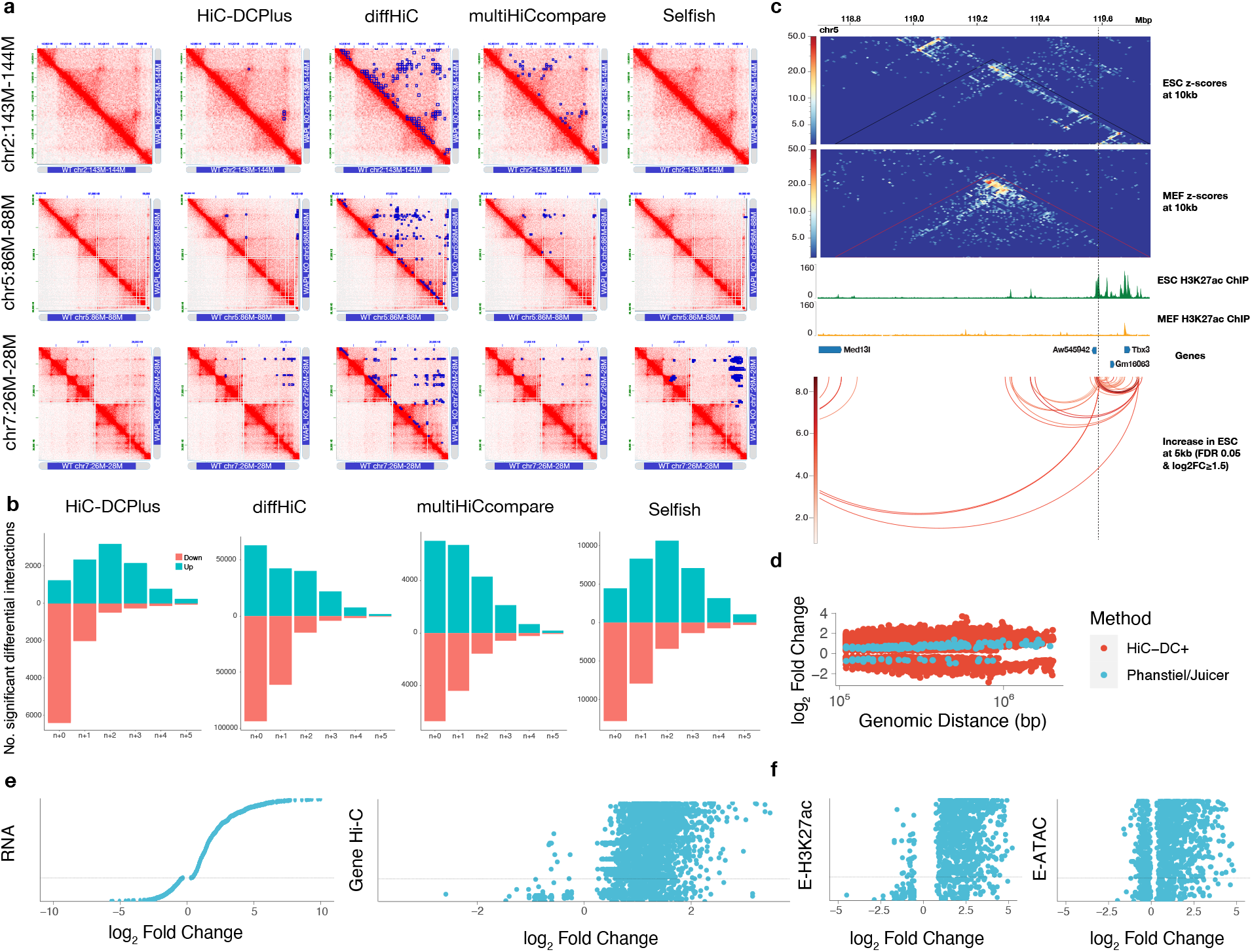
HiC-DC+ differential Hi-C and HiChIP analysis validates global chromatin folding changes during cohesin perturbation and cellular differentiation. **a.** ICE-normalized Hi-C counts for HAP1 WT (lower triangle) and HAP1 WAPL knockout (upper triangle) at 10kb resolution were visualized for three genomic regions highlighted in Haarhuis et al., 2017^17^. For each of these regions, differential Hi-C interactions between *ΔWAPL* knockout vs. WT cells identified by each method at 25kb resolution are shown as blue squares. **b.** Differential interaction analysis between each TAD and its five flanking TADs. The barplots show the number of differential interactions in *ΔWAPL* vs wild type belonging to each category as identified by each method at FDR < 0.05. **c.** HiC-DC+ mESC-specific H3K27ac HiChIP interactions recover the *Tbx3* enhancer hub with *Aw545942, Gm16063*, and *Tbx3* genes at 5kb resolution, all validated targets from Di Giammartino et al., 2019. **d.** Distribution of log fold changes of statistically significant looping events identified by HiC-DC+ and modified Juicer (Phanstiel et al., 2017) as a function of genomic distance (log scale). **e.** Log fold changes of Hi-C interactions and gene expression between THP-1 macrophages vs. monocytes. Each dot in the Hi-C plot represents an interaction between the promoter of a differentially expressed gene and a putative enhancer (**Methods**), while each dot in the RNA-seq plot represents a differentially expressed gene. **f.** Log fold changes of H3K27ac ChIP-seq and ATAC-seq signal between THP-1 macrophages vs. monocytes at enhancer anchors of the differential promoter-enhancer loops identified by HiC-DC+ at 5kb resolution. Each dot represents a putative enhancer.

We also performed contact frequency analysis between each TAD and its five flanking TADs to assess whether differential interaction analysis can reveal TAD aggregation upon WAPL deficiency. We plotted the number of differential interactions identified by each method at FDR < 0.05 that are lost or gained in *ΔWAPL* vs wild type (**Fig. 3b**). Haarhuis et al. claimed that WAPL deficiency mediated gains in interactions at the corners of TADs and loss of intra-TAD interactions. Of the methods tested, only HiC-DC+ recovered both findings: a majority of gained interactions connected neighboring TADs together with a drastic loss of intra-TAD interactions.

To evaluate differential HiChIP interactions, we examined H3K27ac HiChIP data from Di Giammartino et al.^18^ This study mapped high-resolution regulatory loops in mouse embryonic fibroblasts (MEF) and murine embryonic stem cells (mESC) and discovered enhancer hubs, defined as sets of highly connected enhancers interacting with cell-type-specific genes. Further, they disrupted KLF binding motifs within mESC enhancer hubs using CRISPR-Cas9 to test the hypothesis that enhancer hubs coordinate the expression of pluripotency-associated genes. Inactivation of enhancer hubs resulted in coordinated downregulation of all connected genes without affecting the neighboring non-hub genes in mESC. We used HiC-DC+ differential HiChIP analysis to identify mESC-specific enhancer hubs that are lost in MEF. HiC-DC+ detected mESC-specific interactions of the *Tbx3* enhancer hub with *Aw545942, Gm16063*, and *Tbx3* genes, all validated target genes; consistent with validation experiments, HiC-DC+ did not find a differential interaction with *Med13l*, whose expression is not affected by inactivation of this hub (**Fig. 3c**). HiC-DC+ also captured increased interaction of *Zfp42* enhancer hub with *Zfp42* and *Triml2* genes in mESC (**Supplementary Fig. 7**). Notably, another differential HiChIP interaction tool, diffloop^19^, could not detect any differential interactions comprising these two hubs.

Finally, to demonstrate the value of HiC-DC+’s rigorous differential analysis, we reanalyzed very high resolution in situ Hi-C data from the THP-1 cell line model of monocyte to macrophage differentiation, together with epigenomic and transcriptomic data sets to relate chromatin dynamics to gene expression changes^9^.

The original study used Juicer (HiCCUPS) to call interactions and adapted DESeq2 for differential interaction analysis but identified only 217 differential looping events genome-wide at a 10 kb resolution (nominal *P* < 0.001). This modest number led the authors to propose a model where macrophage-specific genes are upregulated through activation of preexisting DNA loops together with a smaller number of gained loops. By contrast, HiC-DC+ identified 64,844 differential looping events at 10 kb resolution (adjusted *P* < 0.01). Of the original differential loop calls, 169 were also called by HiC-DC+ (FDR < 0.01) and 147 of these were found to be differential (adjusted *P* < 0.01). Indeed, we found that HiC-DC+ identified more differential looping events at every genomic distance, and these differential loops exhibited stronger effect sizes than those in the original study (**Fig. 3d**). These results suggest that Juicer is not only too conservative in calling loops but also preferentially identifies loops that exhibit little dynamic behavior. Therefore, we revisited the association of regulatory loop dynamics and gene expression changes.

We reran HiC-DC+ at 5 kb resolution in order to more precisely identify potentially regulatory interactions and found 25,691 differential looping events (adjusted *P* < 0.05). We next considered differentially expressed genes between THP-1 monocyte and macrophages (FDR < 0.05) and found that 1673 (out of 6,782) had at least one promoter-anchored differential loop. Strikingly upregulated genes were associated with gained/strengthened promoter-anchored loops, while downregulated genes were associated with lost/weakened promoter-anchored loops. Moreover, differential promoter-anchored loops at upregulated genes tended to be shorter range interactions (median genomic distance 135kb), while those at downregulated genes were longer range (median genomic distance 185kb, **Supplementary Fig. 8a, b)**. GO enrichment for genes with differential promoter-anchored loops identified functional annotations associated to monocyte to macrophage differentiation (**Supplementary Fig. 9**). These results all suggest that gain or loss of promoter-anchored loops lead respectively to up- or downregulation of gene expression in monocyte to macrophage differentiation, presumably through differential connections to enhancers.

To investigate whether differential promoter-anchored loops indeed include promoter-enhancer interactions, we next constructed an atlas of enhancers using ATAC-seq peaks that overlap with H3K27ac ChIP-seq peaks as a proxy for active enhancer elements (**Methods**). We then restricted analysis to looping events between enhancers in the atlas and promoters (**Fig. 3e**). Upregulated genes were associated with gained/strengthened promoter-enhancer loops, while almost all lost/weakened promoter-enhancer loops were associated with genes with no significant expression change or those that were significantly downregulated. Further, H3K27ac ChIP-seq and ATAC-seq changes at the interacting enhancer elements correlated with expression changes of the target gene, while weaker changes in ATAC and H3K27ac signal were observed at the promoter (**Fig. 3e,f**, **Supplementary Fig. 10)**. Interestingly, CTCF changes at those promoter anchors were strongly associated with the changes in gene expression, although the enhancer sites displayed at best weakly concordant changes in ATAC-seq or CTCF ChIP-seq signal (**Supplementary Fig. 10**). Phanstiel et al. identified multiple gained loops at the *MAFB* locus, and HiC-DC+ analysis also found many enhancer-promoter interactions for *MAFB* (**Fig. 1c**). We also recovered additional genes with previously unidentified differential looping events such as at *GALNT4*, a gene involved in the mucin glycosylation pathway (**Supplementary Fig. 11**).

We have shown that HiC-DC+ facilitates genome-wide analyses of Hi-C and HiChIP data sets and integration with 1D epigenomic and transcriptomic data, recovering previous biological findings in a systematic and global fashion while enabling new insights on the role of 3D genomic architecture in gene regulation. Therefore, HiC-DC+ addresses an important gap in currently available tools for 3D interaction calls and differential analysis and can enable advances in genome-wide understanding of chromatin architecture and dynamics.

## Methods

### Pre-processing of Hi-C and HiChIP data

We aligned HiChIP reads to hg19 or mm10 genomes and filtered out reads that are duplicate or invalid ligation products using the HiC-Pro pipeline (v_2.11.1) with default settings. We processed Hi-C reads using Juicer pipeline (v_1.7.6) with default settings.

### TAD annotations

We used sub-TAD annotations from Arrowhead for GM12878 Hi-C data (GSE63525). We found TADs at 50 kb resolution using TopDom (v_0.0.2) with “w” as 10 on ICE normalized Hi-C counts.

### Calling loops using FitHiChIP, MAPS, and hichipper

We used 5kb-binned hichipper interactions in GM12878 SMC1a and K562 H3K27ac HiChIP reported by ^11^ (GM12878 SMC1a from Table_GC-ALL table, K562 H3K27ac from Table_KH-ALL table). MAPS and FitHiChIP (L) interactions (FDR < 0.01) were also obtained from ^11^ (GM12878 SMC1a from Table_GC-ALL table) and interactions, called using pooled replicates, up to 1.5M were used for comparisons.

We ran MAPS for H3K27ac HiChIP in K562 data with HiChIP inferred ChIP peaks (Table Table_KH-L-H from ^11^, with the following parameters: “bin_size = 5000; fdr = 0; filter_file = None; generate_hic = 0; mapq = 30; length_cutoff = 1000; threads = 10; per_chr = True, binning_range=1500000” using pooled allValidPairs file.

We ran FitHiChIP on pooled H3K27ac HiChIP in K562 data with HiChIP inferred ChIP peaks (Table Table_KH-L-H from ^11^), with the following parameters: IntType=3; BINSIZE=5000; LowDistThr=5000; UppDistThr=1500000; UseP2PBackgrnd=0; BiasType=1; MergeInt=1; QVALUE=0.01” by providing pooled allValidPairs file.

### Benchmarking with Flow-FISH results

We use the 5,091 candidate enhancer-promoter-target gene data from Fulco et al.^12^ reported in Supplementary Table 6a for K562 cells and add 15 singleton experiments for candidate enhancer-promoter pairs along with their tested target genes^20^ for K562 cells reported in Table S3A. Because other methods (all except ABC and HiC-DC+) do not check for interactions shorter than 5kb (17 candidates) and HiC-DC+ longer than 1.5 Mb (324 candidates) for K562 H3K27ac HiChIP, we removed these candidates from consideration. Among those within the distance range, we also removed candidate elements on gene promoters (1,147 candidates) following Fulco et al.^12^ This leaves a set of 3,618 candidate enhancer-promoter-target gene experiments. We treated enhancer-promoter pairs reported at FDR < 0.05 as significant enhancer-promoter candidates for the tested target gene.

### Effect size comparison

We collected significant (FDR < 0.01) looping events identified in K562 H3K27ac HiChIP at 5kb bins overlapping with candidate enhancer-promoter pairs for each of the benchmarked methods (PET count >=2 for hichipper) and compared the reported fraction change in gene expression using one-sided Wilcoxon rank sum test with the alternative hypothesis being HiC-DC+ having greater reduction in gene expression on the CRISPR-perturbed candidates.

### Gene-wise performance comparison

We grouped 3,618 candidate enhancer-promoter pairs by their 71 target genes, chose 22 target genes (2,609 candidates) that has at least one significant pair and one insignificant pair, and computed the area under the precision-recall curve (auPR) for each target gene and for each benchmarked method using significance scores collected from overlapping 5kb bins for each candidate. We ordered the precision-recall curve in an increasing order of *P* values for HiC-DC+, FitHiChIP, and MAPS, and in decreasing order of PET counts for hichipper. We compared the range of auPR values across methods using one-sided Wilcoxon signed rank tests with the alternative hypothesis being HiC-DC+ having greater auPR. For the comparison between ABD and ABC methods (Wilcoxon signed rank test, *P* = 0.39, N=23, alternative: ABD is greater), we included candidate enhancer-promoter pairs closer than 5 kb as well.

### Finding CTCF motif orientation

We found the CTCF motif orientation of the significant GM12878 SMC1a HiChIP loop anchors (FDR < 0.01, 20kb ≤ loop distance ≤ 1.5M) identified by each method using Juicer tool command MotifFinder (v_1.7.6). We used IDR-thresholded GM12878 CTCF peaks available from ENCODE (ENCFF710VEH).

### SubTAD metaplot generation

For each method, we found the number of significant GM12878 SMC1a HiChIP interactions (FDR < 0.01, 20kb ≤ loop distance ≤ 1.5M) a 5kb bin makes with other regions. Then, using deepTools (v_3.1.1) computeMatrix “scale-regions” and plotProfile, we generated a metaplot of significant interactions over GM12878 subTADs (GSE63525).

### Statistical modeling of interaction bin count data

HiC-DC+ follows the feature selections and model specifications outlined in Carty et al.^1^ with the choice of a negative binomial (NB) model for the read counts rather than a zero-truncated negative binomial distribution. Specifically, we used a GLM based on negative binomial regression to model the Hi-C read counts, and we used the fitted model to estimate the statistical significance (*P* value) to the Hi-C interaction bin counts. Following previous notation^1^, we let **Y** = (*y_ij_*) represent the Hi-C contact map of intra-chromosomal interactions, where *i* and *j* are a pair of genomic intervals (either through uniform binning of the genome or fixed size bins of consecutive restriction enzyme fragments) and the tuple (*i,j*) defines an interaction bin. Each bin has an associated vector of covariates, which we denote as **X** = (*x^dist^*; *x^gc^*; *x^map^*), and *y_ij_* is modelled as a random variable that follows a negative binomial distribution. The regression model is defined as: **P**(**Y**=*y_ij_*|**X**)=*f*(*k*; *μ_ij_*; *α*), where the distribution *f*(*k*; *μ_ij_*; *α*) is a negative binomial distribution with dispersion parameter *α*. The negative binomial mean parameter *μ_ij_* is described with a log-linear model:

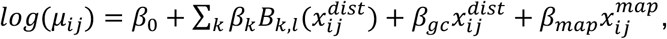

where *β*_0_ is the intercept term, and *β_gc_* and *β_map_* are coefficients for GC content and mappability features, respectively. We modelled the relationship between genomic distance and contact significance with a third order B-spline, which takes the distance covariate as input, and the B-spline basis functions are as previously described^1^. Specifically, we use six degrees of freedom for fitting the spline and have three inner knots. These inner knots are at the 25%, 50% and 75% quantiles of the linear genomic distance.

To increase the statistical power of the model, we removed bins with counts that exceed the 97.5% percentile of null distribution that we deem as positive outliers, corresponding to potentially non-random contacts, and then refit the model to the remainder of the data. We use the function glm.nb in the R library *MASS* to fit the negative binomial regression model.

HiC-DC+ assesses the significance of bin counts based on the corresponding estimated NB distribution for that bin. The *P* value associated with each interaction bin becomes 1 minus the cumulative distribution function fit for that bin given its feature values, i.e.,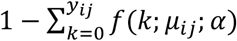.

Significant bins can then be selected on the basis of adjusted *P* values following the Benjamini-Hochberg procedure to control for FDR.

The R package *HiCDCPlus* allows for model selection for other interaction bin count data by providing an option to choose the zero-truncated negative binomial distribution, as well as optional parameters to (i) add/remove/change local genomic features as desired and (ii) use the logarithm of genomic distance as the distance covariate instead of using B-splines.

### Differential interaction calling

We call differential interactions using DESeq2 and raw counts, with two modifications. First, analyzed loci are the interaction bins in the union set of significant interactions (FDR < 0.1) across all conditions to be pairwise tested. Second, size factors for these loci are determined for each genomic distance separately. Following DESeq2 assumptions, we enforce that the median normalized count for interaction bins at each genomic distance should be similar across conditions. Specifically, for a given distance |*i − j*| = *d*, the size factor for condition *k* is chosen to be 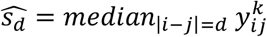, so that normalized counts become 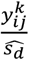.

### Calling differential interactions using diffloop, multiHiCcompare, Selfish, diffHiC

For H3K27ac HiChIP data in mESC and MEF^18^, we called differential loops at 5kb using diffloop 1.16.0^19^ using R version 3.6. Since the published interactions were called in approach similar to Mango^21^, we applied implemented the Mango correction function to remove biases associated with the method. The loops were then filtered at FDR < 0.05 and absolute fold change > 1.5. We performed the differential contact calling on Hi-C data in HAP1 and WAPL knockout HAP1 cells at 25 kb resolution using mutliHiCcompare^15^, Selfish^16^, and diffHiC^14^. All differential interactions were filtered at FDR < 0.05 and absolute fold change > 1.5. After loading raw matrices into multiHiCcompare 1.6.0, we normalized samples with cyclic loess normalization and called the differential interactions by exact test. The python3 version of Selfish 1.10.2 was used to detect differential interactions by applying the method to HiC-DC+ (O/E) normalized matrices. For diffHiC analysis, we loaded raw contact count data as InteractionSet objects, filtered by average counts, used loess normalization, and applied the tool’s statistical model function to estimate *P* values.

### ChIP-seq analysis

We aligned CTCF, and H3K27ac ChIP-seq reads to hg19 using BWA (v_0.7.17-r1188). Then, we extracted uniquely aligned paired reads using SAMtools (v_1.9) and removed PCR duplicates using Picard tools (v_2.18.16). Peak calling was performed for each individual and pooled replicates of each cell type using MACS2 (v_2.1.2) with parameters ‘-g hs -p 0.01’. To find reproducible peaks across replicates for each histone mark, we calculated the irreproducible discovery rate (IDR) using IDR (v_2.0.3) with parameters ‘--samples rep1.narrowPeak rep2.narrowPeak --peak-list pooled.narrowPeak -o --plot’. We combined peaks passing an IDR threshold of 0.05 in each condition for each mark. We used featureCounts (v_1.6.4) to obtain ChIP-seq read counts in the peak atlas, and applied DESeq2 (v_1.24.0) to these counts to find the differential occupancy of each mark between conditions. Bedtools genomeCoverageBed (v_2.27.1) was used to generate bedgraph files scaled with DESeq2 sample size factors and bedgraph files were converted to bigwig using UCSC bedgraph2bigwig (v_4).

### ATAC-seq analysis

We trimmed the raw reads, and filtered for quality using cutadapt (v_2.3). Trimmed reads were aligned to the hg19 genome using Bowtie2 (v_2.3.4.3) and uniquely mapped reads were retained. After centering the reads on the transposase binding event by shifting all positive-strand reads by 4bp downstream and all negative-strand reads 5bp upstream, peak calling was performed on each replicate and all replicated pooled together using MACS2 (v_2.1.2) with parameters ‘-g hs --nomodel’. We calculated the IDR using IDR (v_2.0.3) to find reproducible peaks across replicates. We combined peaks passing an IDR threshold of 0.05 to generate an atlas of peaks. Differential accessibility analysis of the peaks and generation of normalized bigwig tracks was performed as described in ChIP-seq analysis.

### Annotation of peaks

We used the GENCODE transcript annotations of the hg19 (v19) genome to define the genomic location of transcription units. ATAC-seq and ChIP-seq peaks were annotated as promoter peaks if they were within 2kb of a transcription start site. Non-promoter peaks were annotated according to the relevant transcript annotation in the following order of priority: intronic, exonic, or intergenic. Intergenic peaks were assigned to the gene whose TSS or 3’ end was closest to the peak. In this way, each peak was unambiguously assigned to one gene.

### RNA-seq and analysis

We conducted differential gene expression analysis using the gene counts tables provided in GEO:GSE96800 using DESeq2 (v_1.24.0).

### Comparison of macrophage to monocyte differential loops

We reported shared looping events by calling HiC-DC+ differential interactions at 10 kb at an FDR of 0.01 and overlapping with interactions reported by Phanstiel et al.^9^, requiring anchors for our interactions to fall within a 10kb flank of previously the reported anchors.

### Candidate enhancers and promoter-anchored loops

To identify candidate regulatory elements, we first assembled the atlas of reproducible H3K27ac ChIP-seq and ATAC-seq peaks in each the two conditions filtered at an IDR threshold of 0.05. We defined candidate enhancers by taking the intersection of genomic intervals containing both an H3K27ac ChIP and an ATAC peak from this atlas. We identified promoter regions as within 2kb of transcription start sites (TSS) of genes annotated by Gencode GRCh37.p13. We identified elements with differential accessibility or activity of candidate enhancers as based on DESeq2 applied to ATAC or H3K27ac read counts, respectively, with adjusted *P* < 0.05. We described candidate enhancers as static if neither signal was significantly differential. We overlapped CTCF peaks called at IDR 0.05 with candidate enhancer regions and similarly defined differential and static CTCF binding events.

We define promoter-anchored loops as loops and those where one of the anchors overlaps a promoter region. Likewise, enhancer-promoter interactions are promoter=anchored loops with one anchor overlapping a promoter region and the other anchor overlapping a candidate enhancer from the atlas.

### GREAT analysis of differential looping events

We used enhancer-promoter interactions as defined above for GREAT analysis. We filtered results for significance (hypergeometric test, adjusted *P* < 0.05) while restricting to GO terms associated with at most 300 genes.

## Data availability

All data sets used in this study are publicly available and summarized in Supplementary Table 1 with accession codes.

## Code availability

The HiCDCPlus R package is available at https://bitbucket.org/leslielab/hicdcplus.

## Author contributions

M.S. and C.S.L. conceived of the project, designed the statistical methods, and wrote the manuscript. M.S. developed and implemented the methods, developed the software package, and performed benchmarking analyses. W.W. performed computational analyses for monocyte to macrophage differentiation and contributed to writing the manuscript. Y.Z. performed benchmarking experiments for differential analysis under the supervision of R.K. K.V.D. performed statistical analyses for HiChIP benchmarking. C.S.L. supervised the research. All authors read and approved the final version of the manuscript.

