## Supplementary Materials for "HiC-DC+: systematic 3D interaction calls and differential analysis for Hi-C and HiChIP"

**Supplementary Note.** We were curious to see how H3K27ac Hi-ChIP based interaction calls performed relative to the “Activity-by-Contact” (ABC) model for predicting promoter-enhancer loops. The ABC model scores the effect of a putative regulatory element on a gene promoter by taking the product of the enhancer’s activity (A), defined as the geometric mean of DNase-seq and H3K27ac ChIP-seq read counts, and the KR normalized Hi-C contact frequency (C) between the element and promoter; the raw score is normalized by the sum of A times C for all accessible elements within 5 Mb of the promoter. We found that the ABC score outperformed HiC-DC+ HiChIP calls when evaluated by auPR per gene (**Supplementary Fig. 1**). Interestingly, an “Activity-by-Distance” (ABD) score, simply defined by product of the inverse genomic distance with normalized activity, significantly outperforms the ABC score by this measure. Since ABD uses neither Hi-C nor Hi-ChIP, these results suggest that the promoter-enhancer interactions identified by CRISPRi-FlowFISH are enriched for shorter-range interactions that are detectable by H3K27ac signal and accessibility alone. Despite this caveat, enhancer screening data provides a useful benchmark data set for comparing HiChIP interaction callers.

**Supplementary Table 1.** Accession codes for the datasets used in the manuscript

| Cell line | Target protein | Assay | Data used | GEO |
|---|---|---|---|---|
| THP-1 | CTCF | ChIP-seq | fastq | GSE96800 |
| PMA-treated THP-1 | CTCF | ChIP-seq | fastq | GSE96800 |
| THP-1 | H3K27ac | ChIP-seq | fastq | GSE96800 |
| PMA-treated THP-1 | H3K27ac | ChIP-seq | fastq | GSE96800 |
| THP-1 | | ATAC-seq | fastq | GSE96800 |
| PMA-treated THP-1 | | ATAC-seq | fastq | GSE96800 |
| THP-1 | | Hi-C | fastq | GSE96800 |
| PMA-treated THP-1 | | Hi-C | fastq | GSE96800 |
| THP-1 | | RNA-seq | gene counts | GSE96800 |
| PMA-treated THP-1 | | RNA-seq | gene counts | GSE96800 |
| GM12878 | SMC1a | HiChIP | fastq | GSE80820 |
| GM12878 | | Hi-C | Arrowhead domainlist | GSE63525 |
| GM12878 | CTCF | ChIP-seq | optimal IDR thresholded peaks | GSM935611 |
| K562 | H3K27ac | HiChIP | fastq | GSE101498 |
| K562 | | DNase-seq | BigWig | GSM816655 |
| K562 | H3K27ac | ChIP-seq | BigWig | GSM733656 |
| mES | H3K27ac | HiChIP | fastq | GSE113339 |
| MEF | H3K27ac | HiChIP | fastq | GSE113339 |
| mES | | Hi-C | fastq | GSE96107 |
| HAP1 | | Hi-C | fastq | GSE74072 |
| WAPL KO HAP1 | | Hi-C | fastq | GSE95521 |

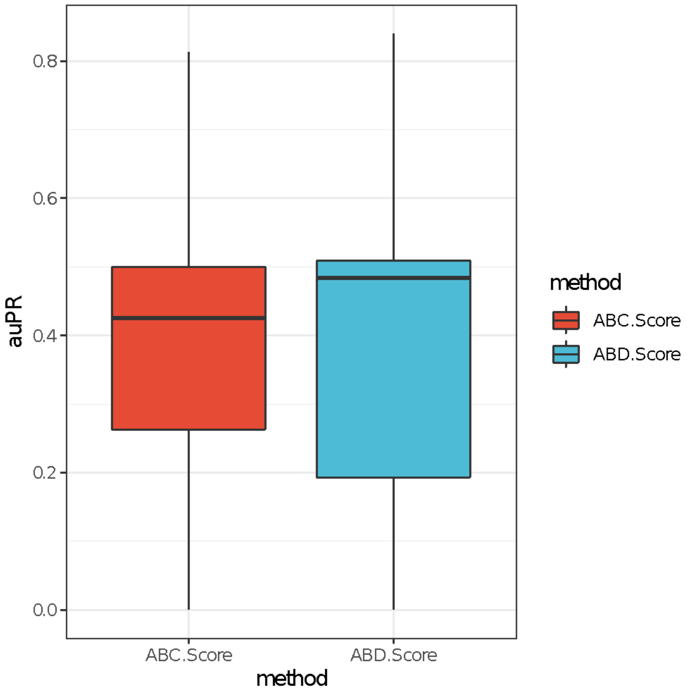


**Supplementary Figure 1.** Comparison of per gene auPR values between Activity-by-Contact and Activity-by-Distance methods using CRISPRi-FlowFISH data in K562 cells (Fulco et al., 2019).

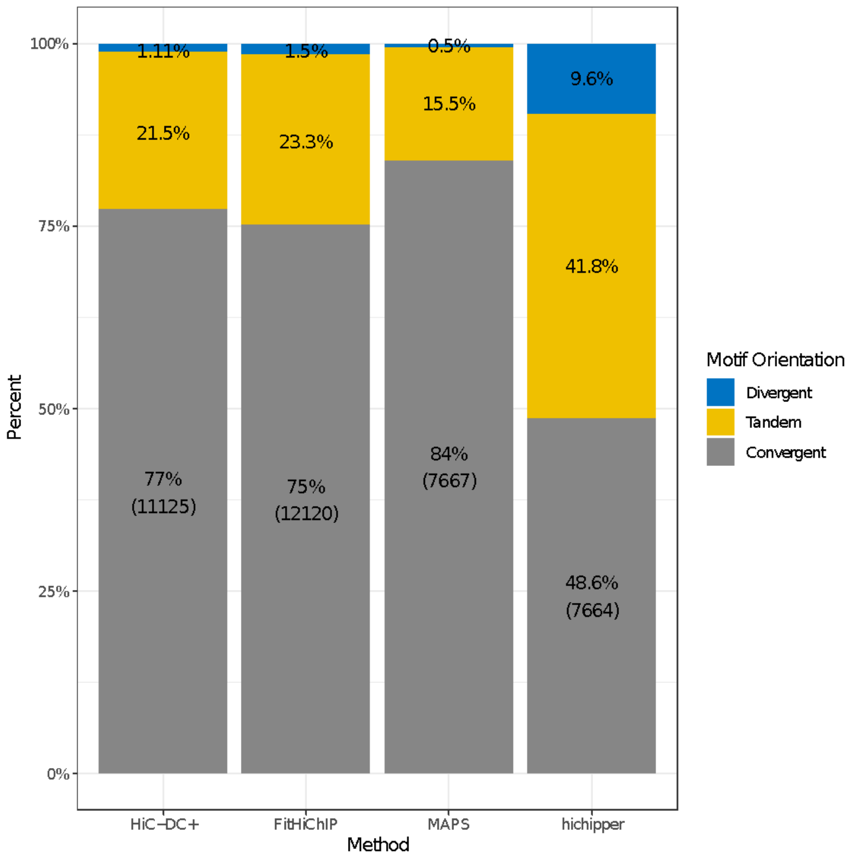


**Supplementary Figure 2.** CTCF motif orientations of GM12878 SMC1a HiChIP loop anchors across different HiChIP interaction callers. We restricted the analysis to loops whose anchors both overlapped with CTCF motifs.

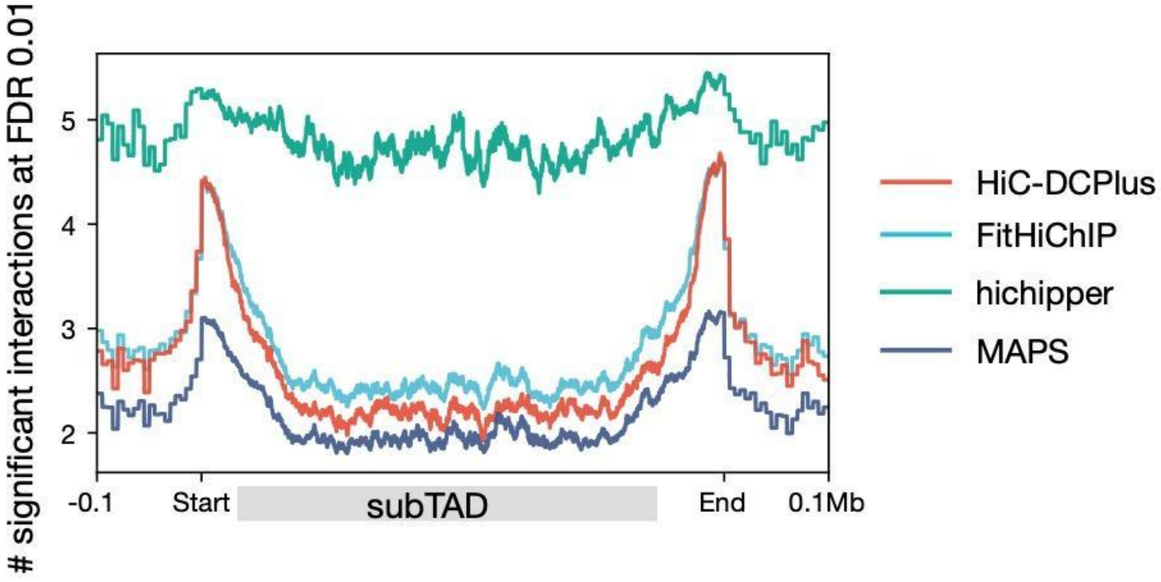


**Supplementary Figure 3.** Metaplot of the number of significant GM12878 SMC1a HiChIP interactions (FDR < 0.01) anchored at each 5kb bin relative to GM12878 subTADs (GSE63525).

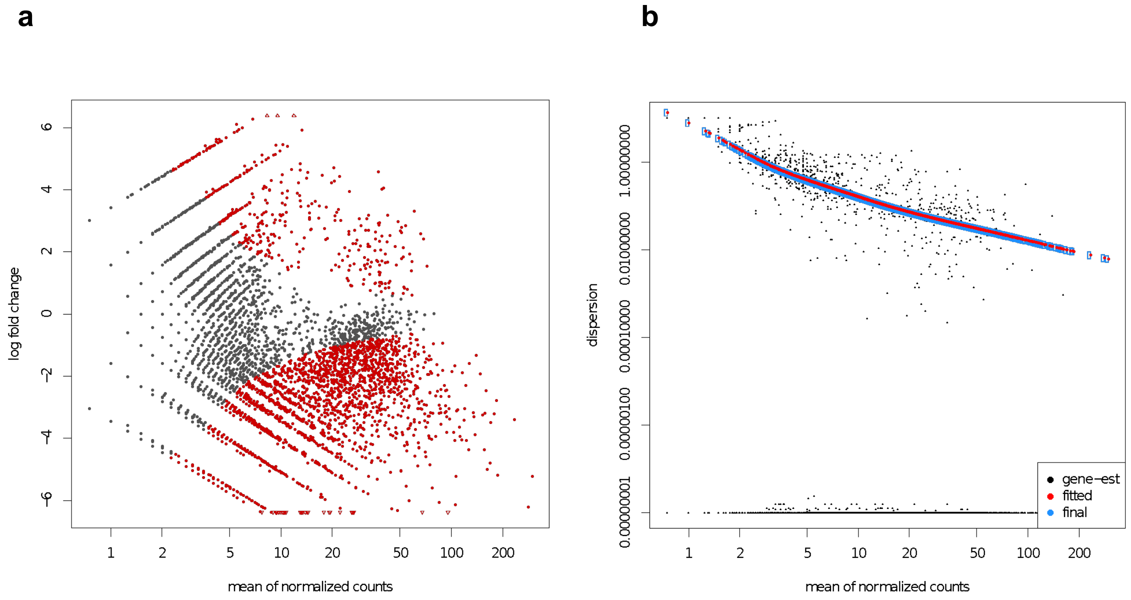


**Supplementary Figure 4.** Diagnostic plots for differential interaction calling. **a.** MA-plot and **b.** Plot of dispersion as a function of mean of normalized counts for differential interaction analysis of H3K27ac HiChIP for mES vs. MEF over chr2 at 5kb resolution.

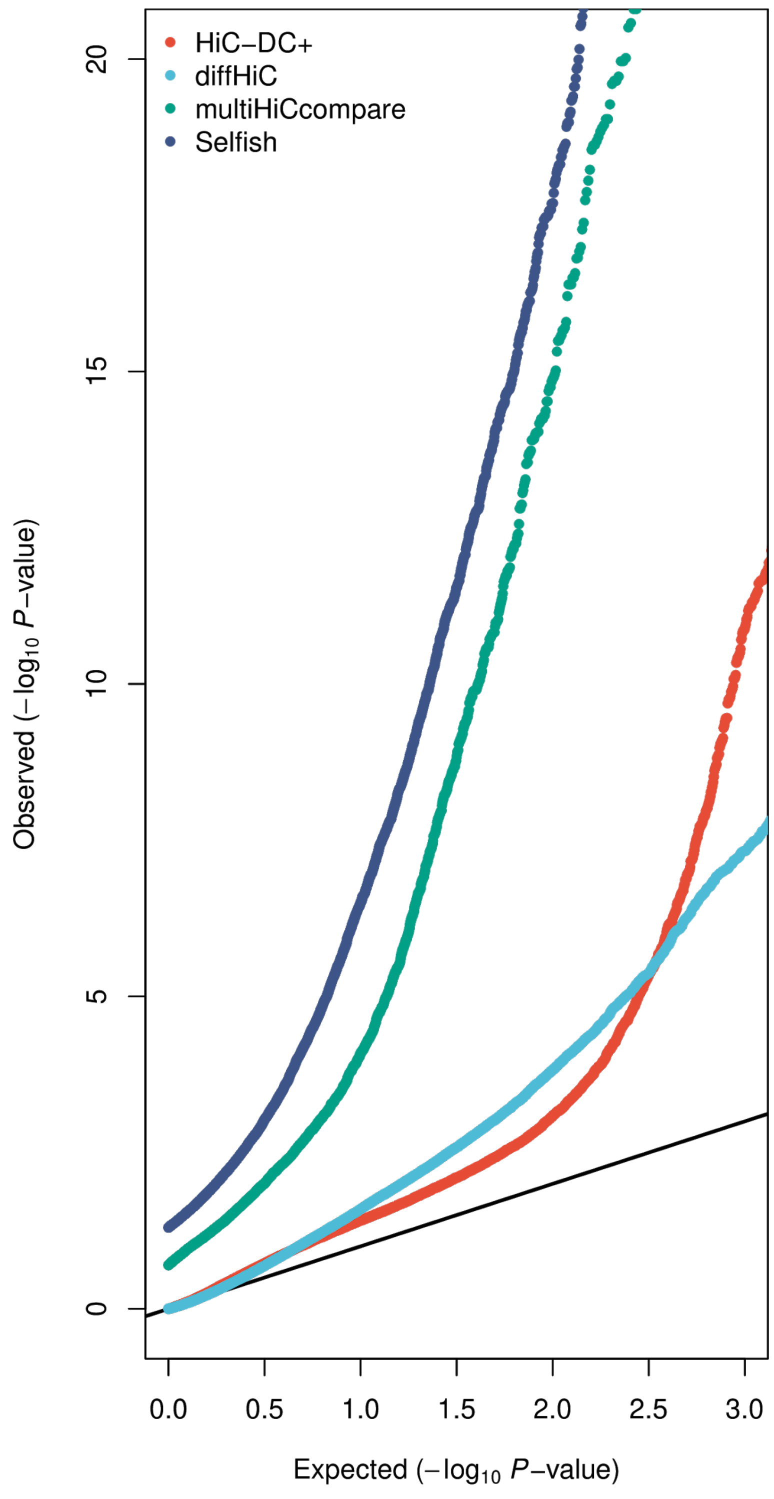


**Supplemental Figure 5.** Q-Q plots of differential Hi-C interaction calling methods on WAPL KO HAP1 vs. HAP1 at 25kb resolution. We restricted this analysis to interactions defined by HiC-DC+ (FDR < 0.1), as HiC-DC+ differential calling was run on this set of interactions instead of the whole genome.

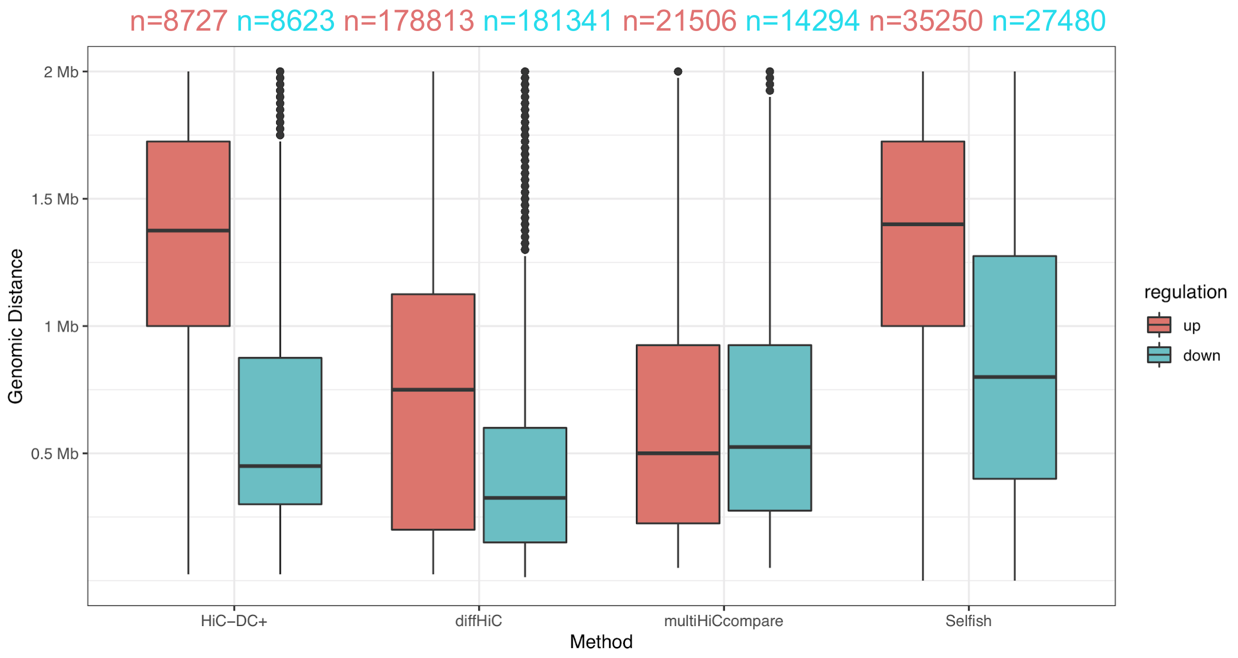


**Supplementary Figure 6.** Loop lengths of differential interactions (FDR < 0.05) called by different methods on WAPL KO HAP1 vs. HAP1 at 25kb resolution. We filtered out differential interactions with distances equal to 0 or more than 2Mb.

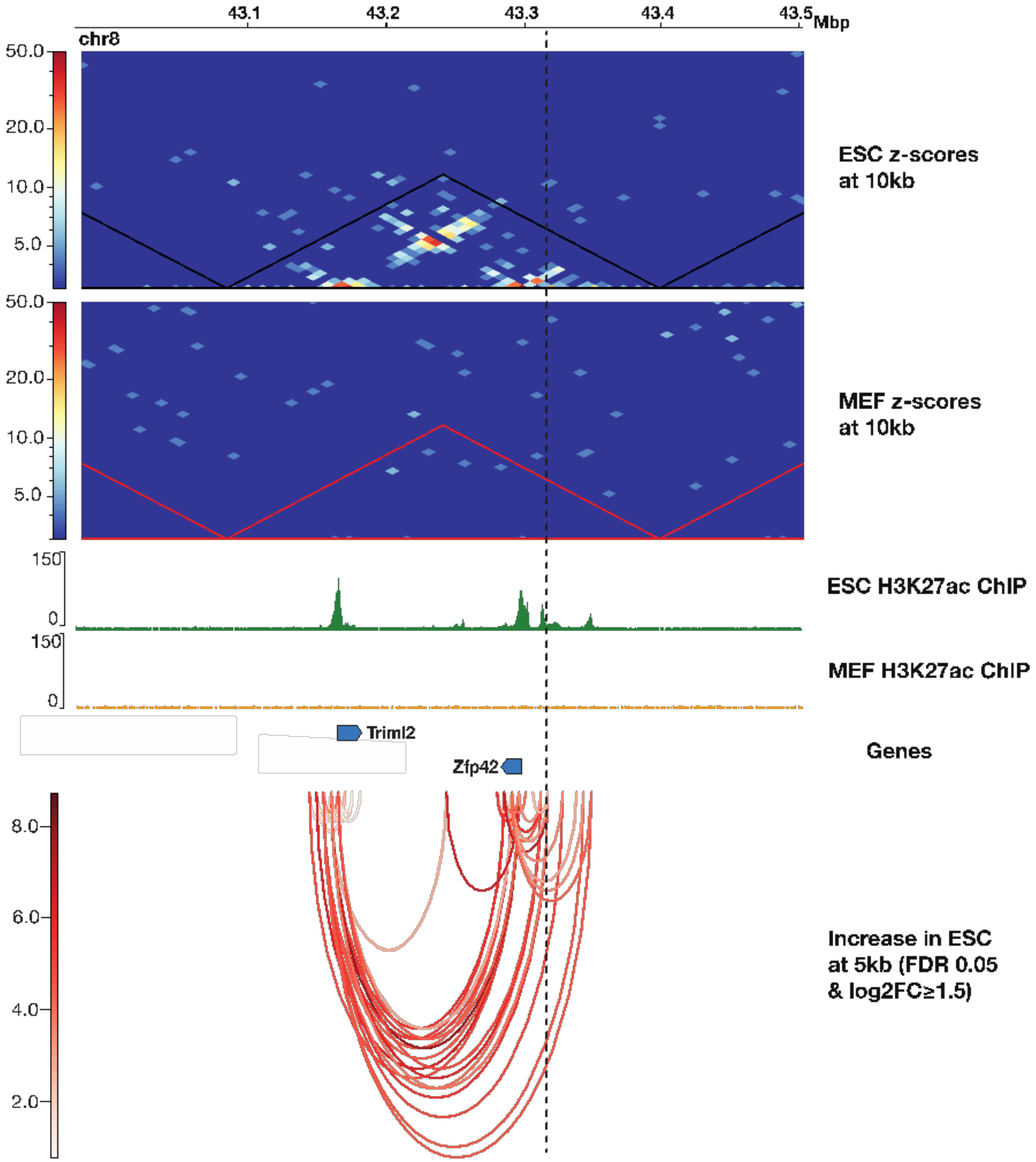


**Supplementary Figure 7.** HiC-DC+ detected mESC-specific H3K27ac HiChIP interactions of an enhancer hub comprising *Zfp42* and *Triml2* at 5kb resolution.

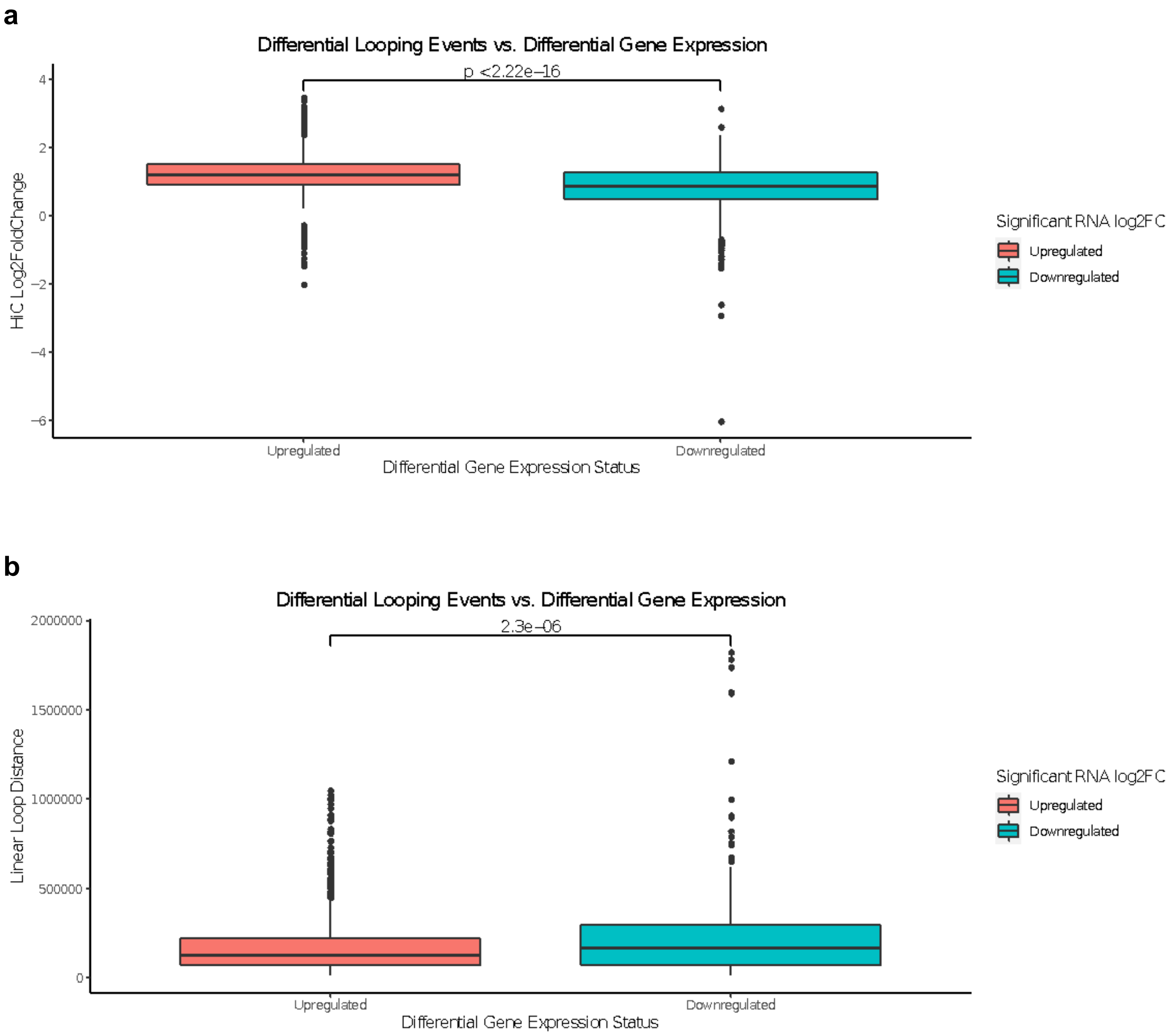


**Supplementary Figure 8.** Distribution of log fold changes (**a**) and distances (**b**) of significant differential Hi-C interactions (FDR < 0.05) with one anchor overlapping the promoter of a differentially expressed gene (FDR < 0.05) between THP-1 monocyte and macrophages at 5kb resolution. Upregulated genes are those whose expression increased in macrophages.

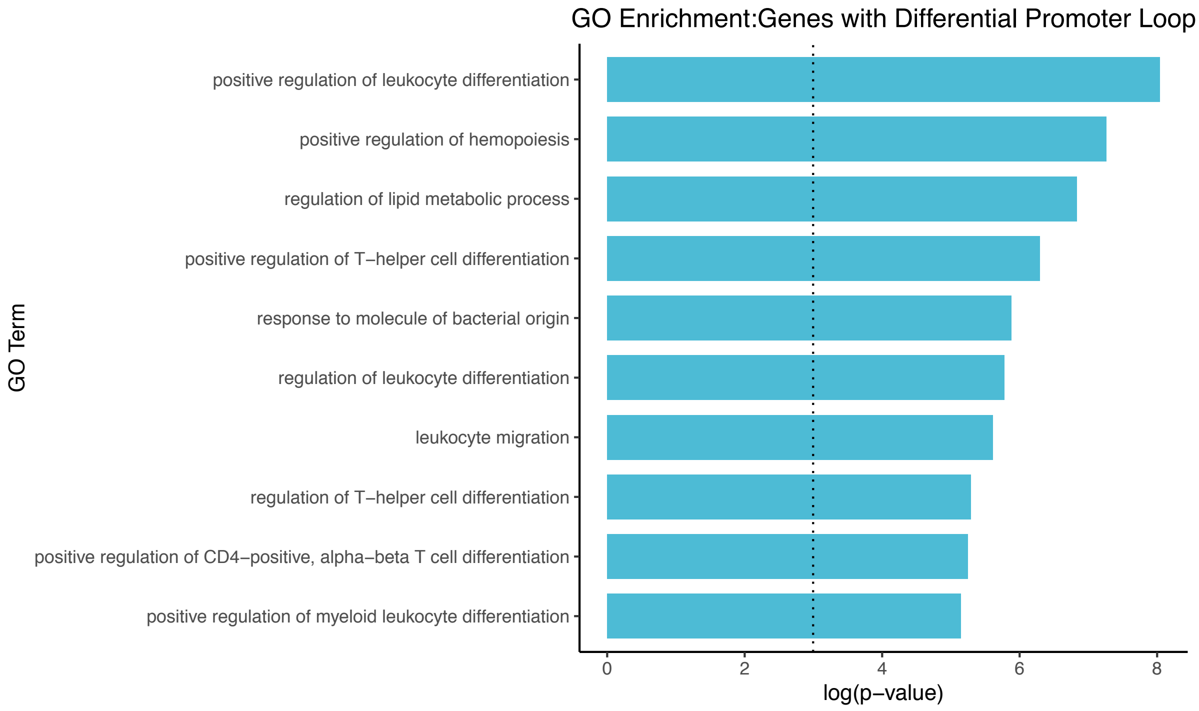


**Supplementary Figure 9.** GO enrichment analysis of genes with differential promoter-anchored loops (FDR < 0.05) in THP-1 macrophages vs. THP-1 monocytes.

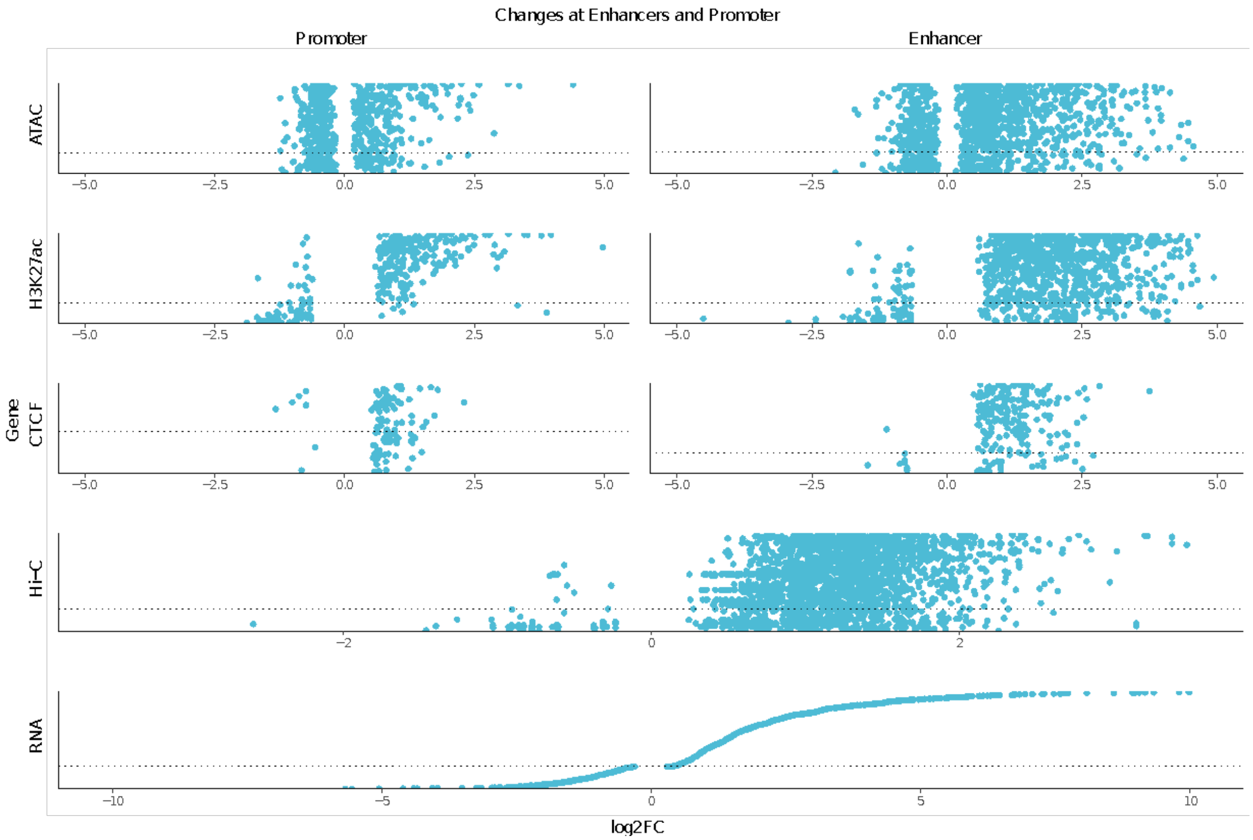


**Supplementary Figure 10.** Changes in ATAC-seq, H3K27ac, and CTCF ChIP-seq at promoter and enhancer anchors of differential promoter-enhancer interactions (**Methods**) between THP-1 macrophages and monocytes at 5kb along with log fold changes of differential Hi-C interactions and corresponding genes.

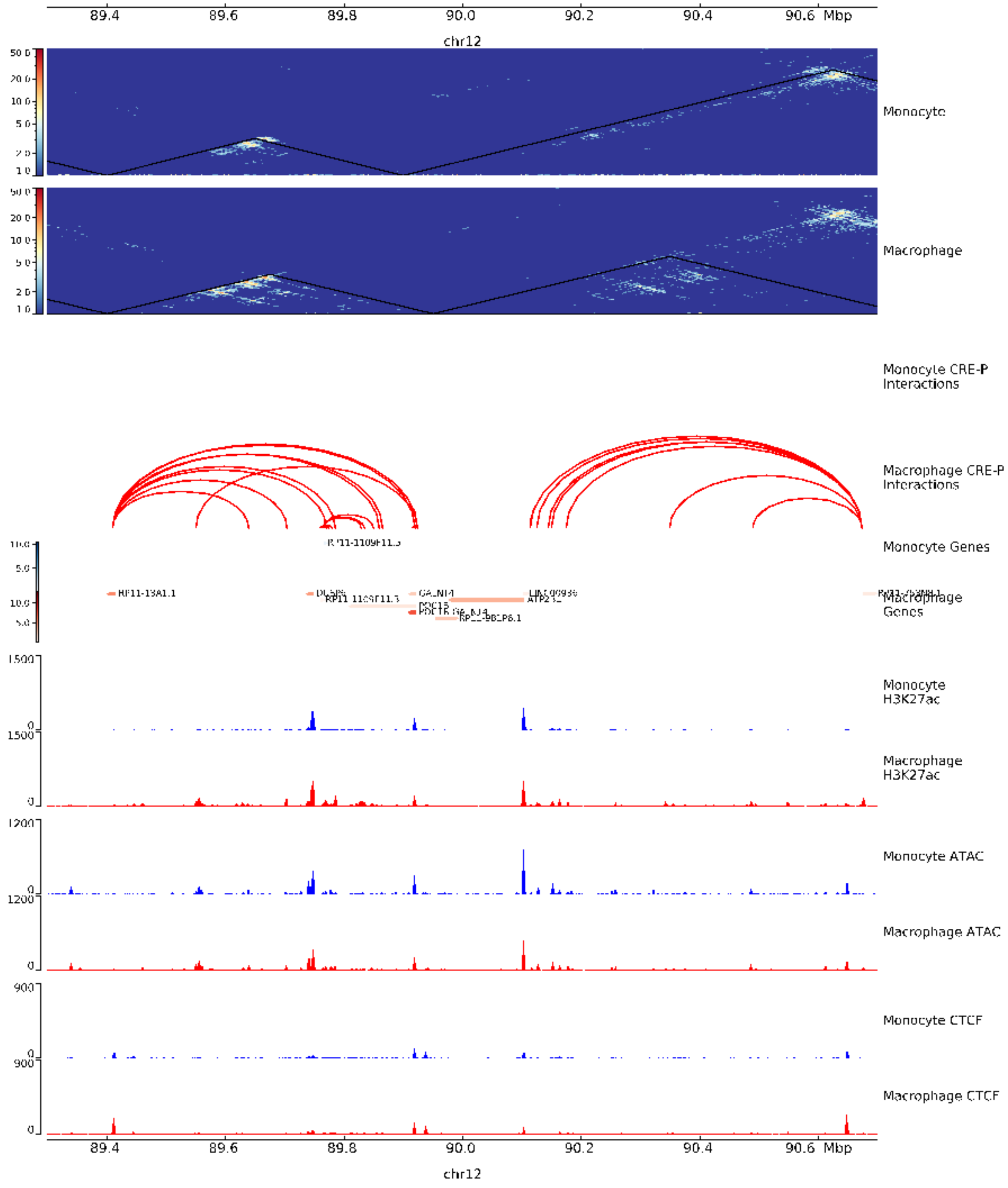


**Supplemental Figure 11.** Differential looping events at 5kb resolution around the *GALNT4* locus along with HiC-DC+ Z-score normalized Hi-C counts (as heatmaps), and CTCF, H3K27ac ChIP-seq and ATAC-seq signals in THP-1 macrophages and monocytes.
